# Soil health pilot study in England: outcomes from an on-farm earthworm survey

**DOI:** 10.1101/405795

**Authors:** Jacqueline L. Stroud

## Abstract

Earthworms are primary candidates for national soil health monitoring as they are ecosystem engineers that benefit both food production and ecosystem services associated with soil security. Supporting farmers to monitor soil health could help to achieve the policy aspiration of sustainable soils by 2030 in England; however, little is known about how to overcome participation barriers, appropriate methodologies (practical, cost-effective, usefulness) or training needs. This paper presents the results from a pilot #60minworms study which mobilised farmers to assess over >1300 ha farmland soils in spring 2018. The results interpretation framework is based on the presence of earthworms from each of the three ecological groups at each observation (20cm^3^pit) and spatially across a field (10 soil pits). Results showed that most fields have basic earthworm biodiversity, but 42 % fields may be at risk of over-cultivation as indicated by absence/rarity of epigeic and/or anecic earthworms; and earthworm counting is not a reliable indicator of earthworm biodiversity. Tillage had a negative impact (*p <* 0.05) on earthworm populations and organic matter management did not mitigate tillage impacts. In terms of farmer participation, Twitter and Farmers Weekly magazine were highly effective channels for recruitment. Direct feedback from participants included excellent scores in trust, value and satisfaction of the protocol (e.g. 100 % would do the test again) and 57 % would use their worm survey results to change their soil management practices. A key training need in terms of earthworm identification skills was reported. The trade-off between data quality, participation rates and fieldwork costs suggests there is potential to streamline the protocol further to #30minuteworms (5 pits), incurring farmer fieldwork costs of approximately £1.48 ha^-1^. At national scales, £14 million pounds across 4.7 M ha^-1^ in fieldwork costs per survey could be saved by farmer participation.

## Introduction

There is now a significant interest in sustainable soil management and policy in England to achieve the Department of Farming and Rural Affairs (DEFRA) aspiration of sustainable soils by 2030. A sustainable arable agricultural system is considered to have both sustainable crop production for food security and a ‘healthy’ soil for soil security. However, there have been few soil surveys to inform both land managers and policy makers about the state of farmland soil health in England to best support evidence-based decision making.

Over the past decade there have been a number of successful public soil surveys in England using earthworm populations including the Open Air Laboratories Soil and Earthworm Survey which included 0.4 % sites in arable fields[1]; the Natural England earthworm surveys which included 1.8 % sites in arable fields[2]; and a school citizen science invertebrate survey (0 % sites in arable fields)[3]. Although earthworms are a primary candidate (out of 183 potential biological indicators) for national soil health monitoring[4], there has been limited farmer participation to date. Mobilising farmers to monitor soil health could be an effective way to improve the national sustainability of soil management. For example, the ‘monitoring effect’ where farms taking part in monitoring activities improve their biodiversity faster than farms not taking part in monitoring[5], fits well with sustainable soil policy aspirations for UK agriculture.

Arable soils typically contain 150 – 350 earthworms per m^2^ and high populations (>400 earthworms per m^2^) are linked to significant benefits in plant productivity, including cash crops such as wheat [6]. There are three ecological functional groups: epigeic earthworms break down surface crop residues and their presence is linked to the breeding season success rates of the song thrush (*Turdus philomelos*), the latter whose populations have rapidly declined in England[7]. Anecic earthworms incorporate surface organic matter into the soil; and support water drainage for plant production[8] and deep crop rooting[9]. UK endogeic earthworm species mix organic and mineral components together to form stable aggregates which benefit spring crop emergence and carbon sequestration[10]. In this way, earthworms support both food production, but also wider ecosystem services associated with soil security. There is no evidence that earthworm biodiversity is constrained in the UK[11], and invasive flatworms which are earthworm predators are largely geographically restricted to Western Scotland and Ireland[12]. Thus, arable soil management is a key factor controlling the relative abundance of these ecological functional groups.

In terms of arable soil management, both epigeic and anecic earthworm species are highly vulnerable to conventional tillage[13], meaning earthworm community structures could be used to indicate over-cultivated soils. Crop establishment practices have been dominated by this intensive mechanical cultivation for decades[14], and this continues to be the principal soil management practice for establishing arable crops in England [15]. It is well known that tillage has an adverse effect on the environmental services provided by soils [16]. Over-cultivation impacts soil biological, physical and chemical properties, for example, causing a decline in surface-feeding earthworms to local extinction levels[13, 17], reduces water stable aggregation which increases the risk of erosion and nutrient losses, and may decrease soil organic carbon levels with implications for climate change[18]. It is unclear as to the extent organic matter management can mitigate the effects of tillage, as the impact of these management activities is subject to local conditions[17].

To date, the use of earthworms in national monitoring schemes has been held back by the absence of a standardised methodology [4]. For example, all three ecological earthworm surveys in England over the past decade have used a different methodology [1-3]. These methods differ from the ISO 23611-1 earthworm assessment method which includes formalin as a vermifuge, precluding its application in citizen science projects. A limitation of the largest international survey of farmland earthworm populations (EU FP7 BioBio) was the skilled labour based protocol and high labour cost (on average 4.8 person days (£3 k) per farm for earthworm fieldwork alone, not including taxonomic identification)[5].

The ultimate aim of monitoring is to cost-effectively convey robust information to those who are expected to use it [19]; essentially the trade-off between data quality, practicability, cost and usefulness. The principal cost of monitoring is labour; for which the UK has the highest person day costs in the EU [5]. Research from the EU FP7 BioBio project indicated significant cost reductions (46 %) could be achieved if farmers could be mobilised to assess their own farms; however, key research areas include how to overcome participation barriers; the development of protocols that require lower technical expertise; identification of training needs and quantifying sampling bias [5]. To date, one small study assessed the usefulness of ‘earthworms’ (numbers and species) for farmland biodiversity assessments to administrators, farmers and consumer groups, with earthworms ranked 5^th^ (out of 6 parameters) by all groups [19].

The aim of the #60minworms pilot study was to support farmers to monitor their own field(s) and generate results that are useful to their soil management decisions. A number of gaps in on-farm earthworm monitoring are addressed, and the state of farmland soils in England as indicated by earthworm populations under different monitoring and interpretation scenarios are presented.

### Methods

The #60minworms pilot study (100 fields target) ran between the 15^th^ March – 30^th^ April 2018 (Fig. 1). There was no need for ethical approval as this was undertaken by volunteers (farmers) on their privately-owned land (farms). Survey booklets were distributed directly (at soil health workshops in March) or following a request and posted to potential participants in order to quantify recruitment and participation channels. All the participants received a report on their earthworm populations and were invited to take part in the Rothamsted #60minworms workshop on the 3^rd^ May 2018. The workshop was based around a ClikaPad audience response system to quantify sampling design bias, method compliance, competence, usefulness and future developments, and afterwards, an earthworm identification class was held (at participants request). The outcomes were adopted to make the new Agricultural and Horticultural Development Board (AHDB) factsheets ‘How to count worms’ freely available as printable leaflets in June 2018, with an initial print run of 2000 copies, distributed at agricultural events such as Cereals (leading technical event for the arable industry with up to 20,000 visitors) and AHDB strategic and monitor farm events (24 sites around the UK) (Supplementary Information (SI) booklet).

**Fig. 1:**
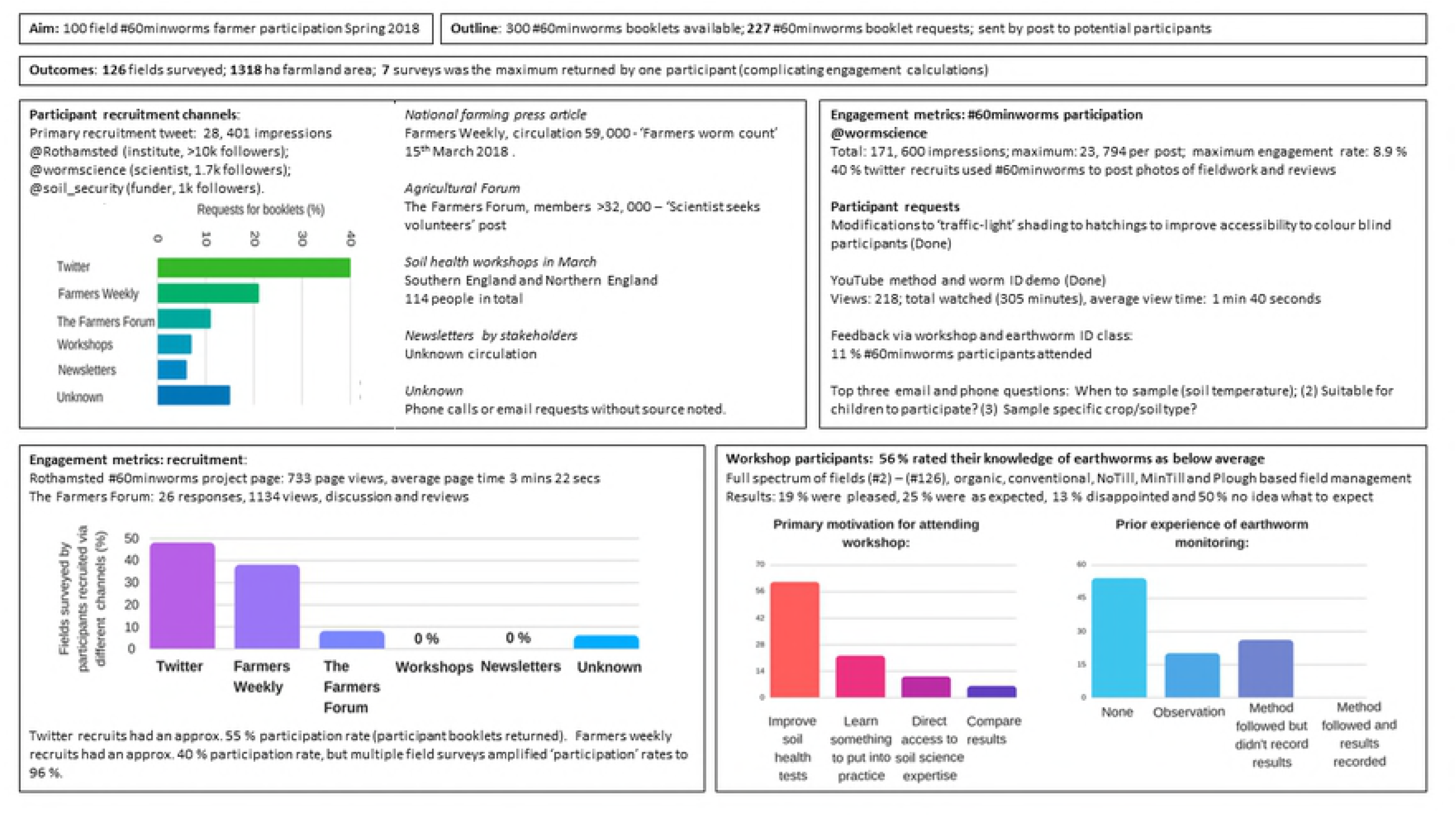
Recruitment, participation and engagement in the #60minworms survey. The key mobilisation routes were through Twitter and Farmers Weekly, the survey attracted participants with no earthworm monitoring experience and the primary feedback preference was a workshop.

The #60minworms methodology was designed around the presence of earthworms in the field, enabling a rapid traffic-light based interpretation. The participants required five pieces of equipment to perform the survey: a garden fork to dig the soil pit, a ruler (as 20 cm size pits needed), a mat (to put the soil on for hand-sorting *in-situ*), a pot with a lid (to stop earthworms escaping) plus a small volume of water (so the earthworms do not dry out) and the results booklet (including a simple earthworm key) with a pen. A timer was recommended to complete the hand-sorting within 5 minutes, unless the soil was too wet or compacted to sort efficiently and time was increased to 10 minutes. Thus, the equipment and consumable costs were negligible; and, an experienced sampler could generally complete the survey in 60 minutes. The procedure was to dig a 20 cm x 20 cm x 20 soil pit and place the soil on the mat. The soil is hand-sorted, placing each earthworm into the pot. Once the soil has been sorted, the total number of earthworms were counted and recorded. The earthworms were separated into adults (for further analysis) and juveniles (returned to the pit). Adult earthworms were separated into an ecological functional group (epigeic, endogeic or anecic) using a simple key. There are high levels of cryptic diversity within UK earthworm species[22], thus species level assessments are beyond the scope of this agricultural soil health assessment. The total numbers of epigeic (small red worms), endogeic (pale or green worms) or anecic (heavily pigmented, large worms) adults were recorded for each pit. After analysis, the adult earthworms were returned to the pit. This was repeated 10 times via a W-style sampling pattern across the cropped field.

To address some of the common concerns relating to earthworm analyses, the seasonal reproducibility was tested on nine AHDB strategic farm fields (eight arable and one grass field) in October 2017 and April 2018. To assess the reliability of 10 or fewer soil pits per field; 20 soil pits per field (n = 9 fields) were measured. To assess the accuracy of hand-sorting earthworms in 5 minutes, sorted soil was re-sorted for 5 minutes and earthworms were collected for further analyses. This was performed by three volunteers on nine fields (range of soil textures and crop types) (n = 27 pit resorted) in April 2018. To indicate year-on-year variability, previous scientific field trial based earthworm surveys[23] (using the identical soil pit size and hand sorting methods), with at least two years of data were re-analysed (to remove vermifuge data and categorise the species into their ecological groupings), and recalculated on a per pit basis using the likelihood formula.

Instant results analysis is possible in five categories: (a) widespread presence, (b) epigeic, (c) endogeic, (d) anecic presence, and (e) presence of earthworm ‘hotspots’ of earthworms via a simple likelihood formula:

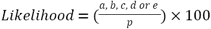

Total number of pits (p) where at least:

a. one earthworm was found (juveniles or any adults below),
b. one adult epigeic earthworm was found,
c. one adult endogeic earthworm was found,
d. one adult anecic earthworm was found,
e. high numbers (≥ 16 earthworms per pit, ≥400 earthworms per m^2^) of earthworms (total number including all juveniles and adults) found

The traffic light system interpretation indicates a red, ‘unlikely’ category (<33 %), the amber, ‘possible’ category (>33 – 66 %) and the green ‘likely’ category (> 66%), and is reported on a field basis (Table 1).

**Table 1:** The interpretation framework is based on the presence of earthworms for each observation (one soil pit) across a field (10 soil pits).

| % Occurrence | Likelihood: |
| --- | --- |
| 0 – 1 | Exceptionally unlikely |
| 1 – 10 | Very unlikely |
| 10 – 33 | Unlikely |
| 33 – 66 | As likely as unlikely |
| 66 – 90 | Likely |
| 90 - 99 | Very likely |
| 99 - 100 | Almost certain |

The satisfactory threshold was the possible ≥40% score for each category, providing evidence for basic earthworm biodiversity in-field; where the likely >66 % score provided good evidence for basic earthworm biodiversity. Scores ≤10 % for presence and/or an ecological group are sub-optimal, as there is little evidence for spatial impacts (widespread presence of an earthworm ecological group to support plant productivity and wider ecosystem services); and temporal impacts (an adults’ lifespan is in the order of years, and given reproduction capacity increases the likelihood of that ecological function being sustained in the future). Earthworm numbers were not of primary interest because the interpretation is dependent on fertiliser usage, soil type, crop type etc[6]. but to calculate the average number of earthworms per hectare the following formula was used:

(f) Earthworms per hectare =(mean number of earthworm per pit × 25) & & × 10000 Whilst the results (simple percentages) could be calculated by the participants, they were requested to either post or email a copy of their findings, and include basic field management details including field name, size, postcode, crop, tillage (notill, minimum tillage and ploughed), and Yes/No answers to organic matter management: residue retained, cover cropping and whether an organic waste e.g. compost had been used this year, in order to inform on general soil management practices and earthworm results. A total of 10 participants with either depleted or exceptional earthworm results were contacted and visited.

Following the submission of all the data, Genstat (18.2.0.18409, 18^th^ addition, VSN International Ltd., UK) was used to perform one-way ANOVAs to assess trends in earthworm populations and soil management practices. Labour cost estimates were calculated using a £:€ exchange rate of 1.12; in order to translate private agency skilled worker (€89.75 h^-1^) and farmer (€28.39 h^-1^) [5]. To calculate costs at farm, regional and national scales, DEFRA official statistics (February 2018) were used [24]. The survey data was compared against the earthworm soil health thresholds proposed in this paper; and the proposed AHDB soil health scorecard earthworm number thresholds[25] to estimate the state of farmland soil health.

## Results

### Recruitment and engagement of farmers

Participants recruited through Twitter had exceptional recruitment and engagement rates of up to 55 %, and engagement was amplified to ‘96 %’ (multiple fields surveyed) by participants recruited through Farmers Weekly (Fig. 1). In contrast, no engagement (0 %) from potential participants recruited via the soil health workshops or newsletters was found. The Rothamsted #60minworms workshop was attended by participants from a diverse range of management practices, primarily interested in improving soil health assessments and no prior experience in earthworm monitoring (Fig. 1).

### Cost and usefulness of the #60minworms survey

Most participants (77 %) reported spending 5-mins hand-sorting each soil pit, enabling completion within 60 minutes. The number of samples was fixed at 10 replicates, but field surveys ranged between 2 to 80 hectares (average observation was 1.08 ± 0.08 pit per hectare) and the longest reported survey took 3 hours. Using the person (farmer) day costs in the UK[5], where the majority (66 %) of participants performed the #60minworms analysis alone means the typical farm labour costs were €28 (£25). A total of 34 % participants completed the survey with fieldwork support provided by up to 3 people, increasing the cost to €84 (£75) per field. The real farm labour costs (in-kind) for the 126 field #60minworm pilot field study can therefore be estimated to be in the order of €5928 (£5300); which on a per hectare basis is €4.50 (£4).

There were a range of motivations for taking part in the #60minworms survey, and excellent scores in value, trust and satisfaction of the method (Fig. 2); for example, 100 % of the participants would do the #60minworms survey again. There were very high scores for community science in every category; where 100 % participants would recommend the survey to others, 93 % of participants rated other participants’ competence was very important and 87 % participants would use of scientific field trials to aid their interpretations; which corroborated with the high (29 %) primary use of results would be to compare their results to others (Fig. 2). Further, most participants would use the survey to compare soil management practices on-farm (36 %); which is in agreement with participants performing multiple field surveys and change their soil management practices based as a result earthworm monitoring results (57 % participants) (Fig. 2). There was no interest in regional trends, with usefulness only linked to relevant comparisons and threshold values (Fig. 2).

**Fig. 2:**
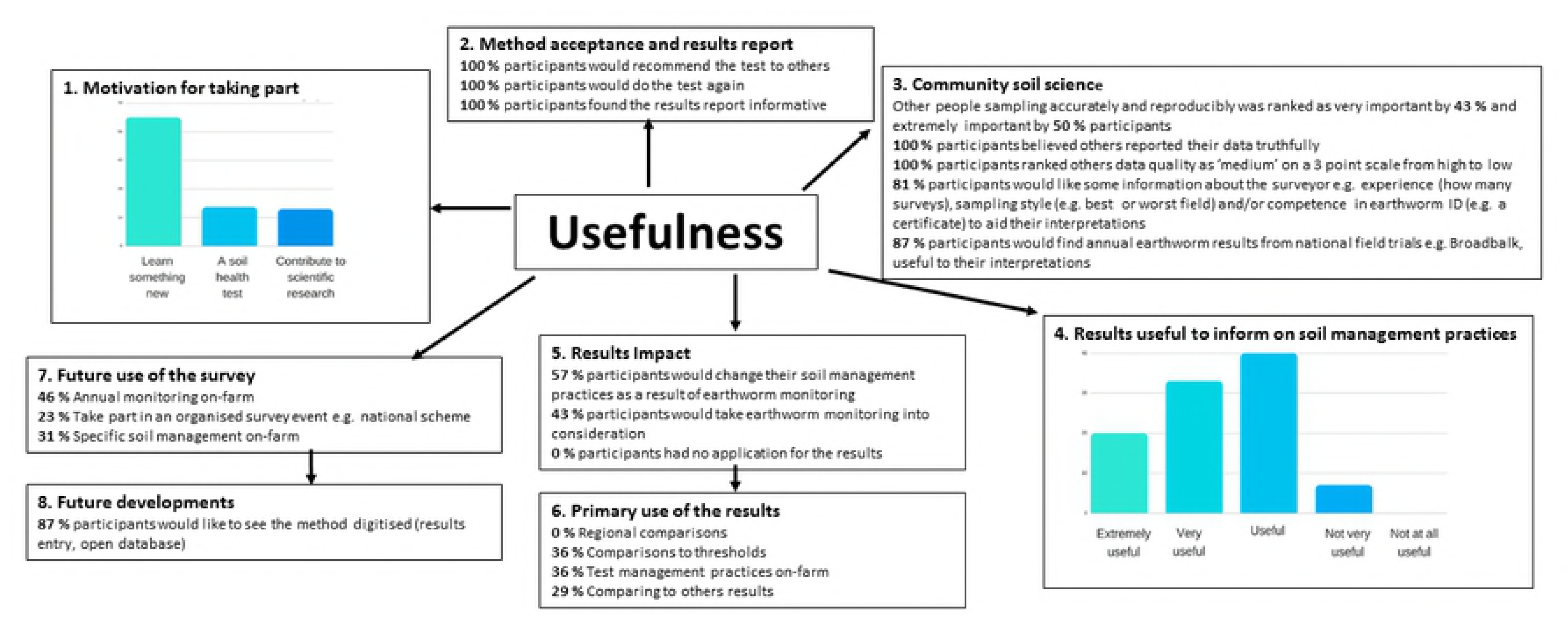
Usefulness of the #60minworms survey to farmers. Feedback included trust, value and satisfaction in the protocol by participants (100 % would do the test again) and an extremely high interest (>85 %) in community science (including other participants and scientists) with a key use in comparing results

### Quality control and application

There was full geographic coverage in England and a range of management practices surveyed (Fig. 3). Choosing the smallest field was not a sampling strategy by any participant, and good levels of compliance were recorded, for example, all participants measured the size of their soil pit(s). A key training need in earthworm identification skills (Fig. 3). Farmers reported a problem capturing deep burrowing *Lumbricus terrestris* anecic earthworms which could be solved by amending the method to include a tick box for the presence of middens/characteristic large vertical burrows. There are three common anecic earthworm species in England (*L.terrestris, A.longa* and *A. nocturna*), and middens are a good indicator of *L.terrestris[26-32]*, the earthworm most sensitive to conventional tillage [13].

**Fig. 3:**
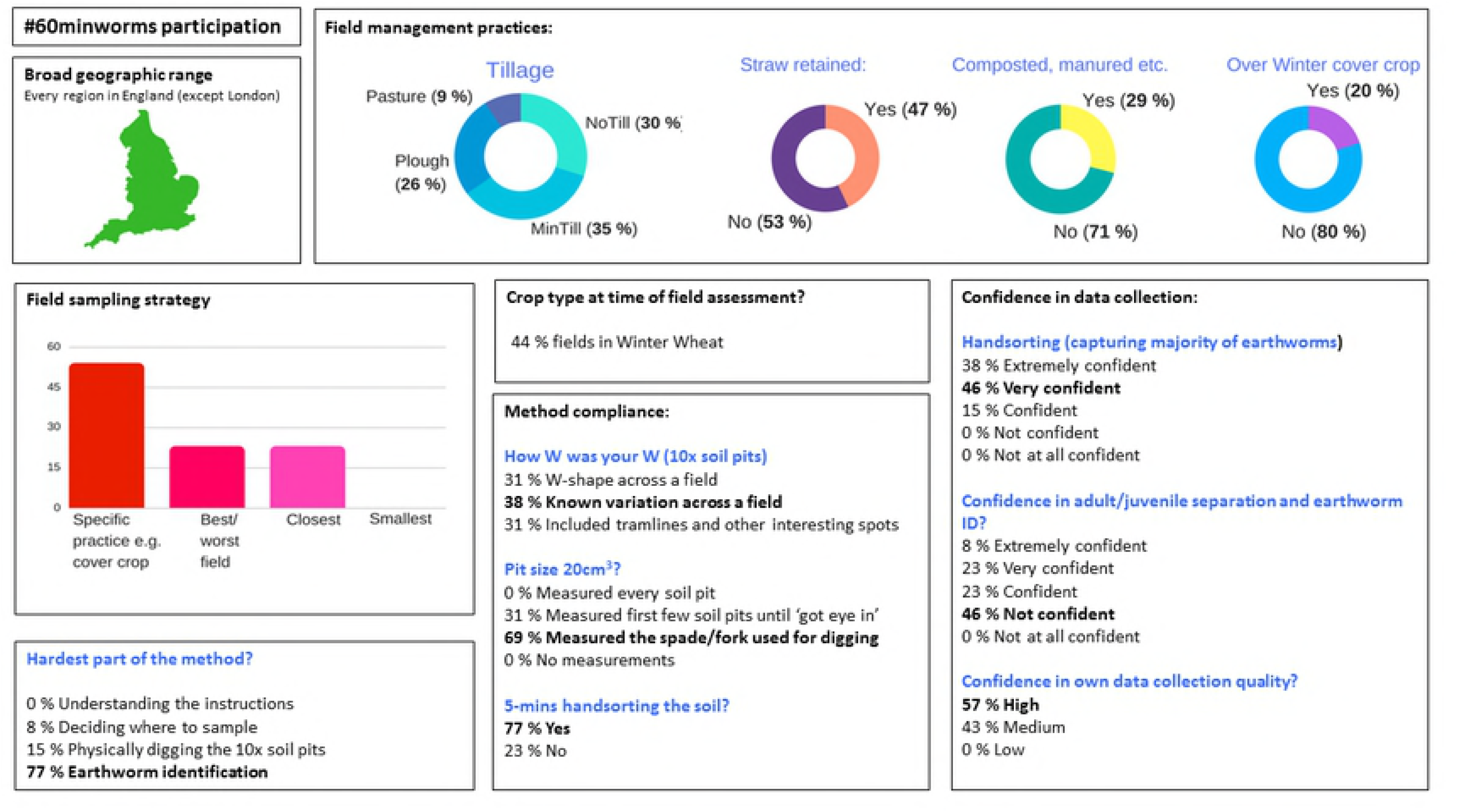
#60minworms survey participation. There was a broad geographic spread over England and a range of field management practices. There was little indication of bias in sampling strategy, problems in compliance or results quality, but there was a key training need in terms of earthworm identification skills.

The intensive sampling at the AHDB strategic farm fields also measured the accuracy of 5- minute soil pit handsorting for earthworms. Resorting soil for a further 5-minutes led to an additional 1.6 ± 0.17 earthworms per pit per field (regardless of earthworm population size), ranging in biomass from 0.05 – 0.429 g per earthworm, of which 91 % were juveniles; meaning the underestimation of 40 worms per m^2^ (or 400, 000 ha^-1^) on each field. The variability of earthworm populations over annual scales was high for earthworm numbers (SI, Table S1); but the presence (or absence) of each ecological group was consistent (SI, Table S1, S2). Comparing results at 20, 10 and 5 sampling pits per field; 10 sampling pits would incur an error of 16 % in categorizing the earthworm groups; of which 4 % would be a false negative (i.e. 0 %, no sightings on that ecological group which is uncommon rather than absent); five sampling pits per field would incur an error of 33% in categorizing the earthworm groups, of which 15 % would be a false negative.

### #60minworms survey results

Earthworm counts within a 10-pit field survey ranged by 6.4-fold, from a minimum 1.3 to a maximum difference of 28-fold. The average earthworm field population was 2.4 ± 0.4 million worms ha^-1^ and ranged by 100-fold, between 0.75 to 7.3 million worms ha^-1^. The field characteristics of the top and lowest 10 populations of earthworms shared soil textures, tillage and field management practices (SI, Table S3). Tillage significantly (*p <* 0.05) impacted the general earthworm presence, epigeic presence, anecic presence, presence of hotspots and number of earthworms per hectare (SI Fig. S1, Table S4). Organic matter management included straw retention, cover cropping or manuring (including animal manures, compost, anaerobic digestate, humic substances or biosolids). The only significant impact on the numbers of earthworms was straw retention (*p* = 0.04), Table S4. Cover cropping, significantly impacted the presence of anecic earthworms (*p* = 0.03), (SI Fig. S2, Table S4).

A total of 77 % fields had a 100 % presence of earthworms (at least 1 earthworm per pit), with the lowest presence recorded at 30 % for one field. There were no sightings of epigeic earthworm on 21 % fields, and anecic earthworms on 16 % fields (Table S5a), with a further 8 – 11 % fields have rare sightings of these groups (10 % presence). There was a good (≥ 67 % presence) of endogeic earthworms on most fields (Table S5a); and a good presence of all three ecological groups together on 15 % fields. Earthworm hotspots (≥16 earthworms per pit) were uncommon; 46 % fields had no earthworm hotspots, where a good presence of hotspots was detected on 13 % fields. Overall, 42 % fields had sub-optimal earthworm populations, defined as ≤10 % presence for at least one ecological group, providing little evidence for the spatial and temporal presence of epigeic, endogeic and/or anecic earthworms.

### Trade-offs between data quality, participation rates and cost

The aim of #60minworms was to indicate soils at risk of over-cultivation through the absence/rarity of epigeic and anecic earthworms that have well known sensitivity to tillage. Reducing the sampling intensity to five soil pits (e.g. #30minworms) and changing the sub- optimal threshold to <20 %, shows good agreement to the 10-pit survey ≤10 % category threshold (Tables S5b, c). An alternative metric is to rate the soil health of a field based on earthworm numbers at a sampling intensity of one soil pit per field as proposed for the AHDB soil scorecard[25]. This survey indicates that between 68 – 88 % fields could be categorized as ‘depleted’ through to ‘active’ (Table S6). In comparison a sampling intensity of five soil pits per field provided average earthworm count data that was in good agreement with these data calculated at 10 soil pits per field (Table S6), and 20 % of fields would be categorized as ‘depleted’ at this sampling intensity. However, even at a high soil pit replication (n = 10) there was a limited concomitant relationship between #60minworms ecological group absence(s) and AHDB soil scorecard ‘depleted’ earthworm numbers in field classifications (Table 2). For example, no adult earthworms were found on six fields; but 33 % of those fields were classified as ‘active’ as earthworm number thresholds are weighted towards juvenile earthworm abundances.

**Table 2:**
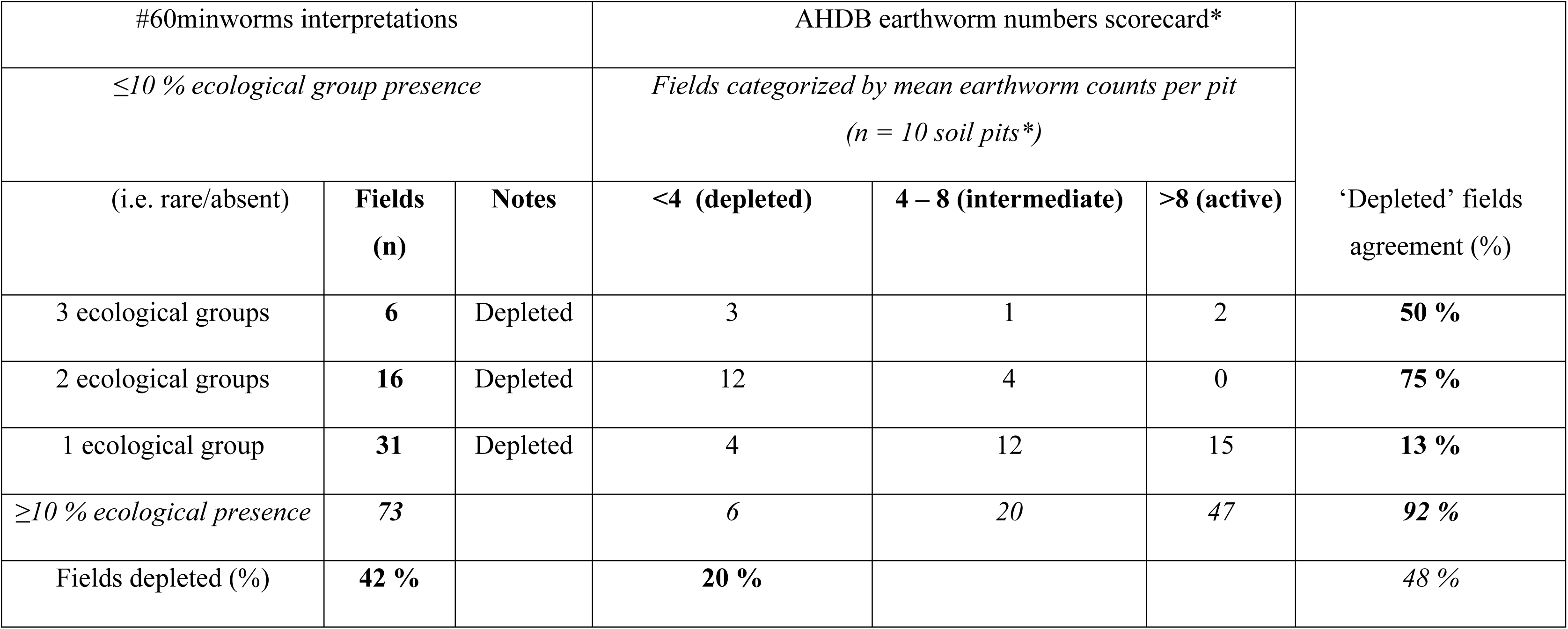
Fields interpreted as ‘depleted’ differ between the interpretation method, noting that the proposed AHDB numbers scorecard analysis is based on the mean of 10 soil pits (*not one pit as proposed for this method). A total of 42 % fields may have over-cultivation issues (no evidence of ecological group presence); where as ‘depleted’ earthworm numbers occurred on 20 % fields; these indicators do not necessarily co- occur.

The trade-off was estimated using data quality (% false negatives), participation (scaling booklet requests to 100 % and actual survey time to 100 %) and cost (using an intensive 20 pits x 10 minutes earthworm fieldwork set at 100 %), indicates that a five-pit field survey has significant potential (Fig. 4). An average #30minworms field survey (10.9 ± 0.8 ha^-1^) would incur £16 – 48 in fieldwork costs depending on labour type (farmer or outsourced). Scaling to #30minworms of the whole arable area (52 %) of an average farm in England (85 ha) would range between £65 – 196 in fieldwork costs depending on labour type. Significant regional variations in farm costs would be expected; fieldwork costs on the arable area on an average farm in the North East being £23 - 70, where the East of England would cost £134 – 401; reflecting farm size and arable cropping area. Nationally, a #30minworms survey of the entire 4.74 million hectares of land under arable cropping would have fieldwork costs at £7 million (farmer participation) to £21 million (outsourced) per survey.

**Fig. 4:**
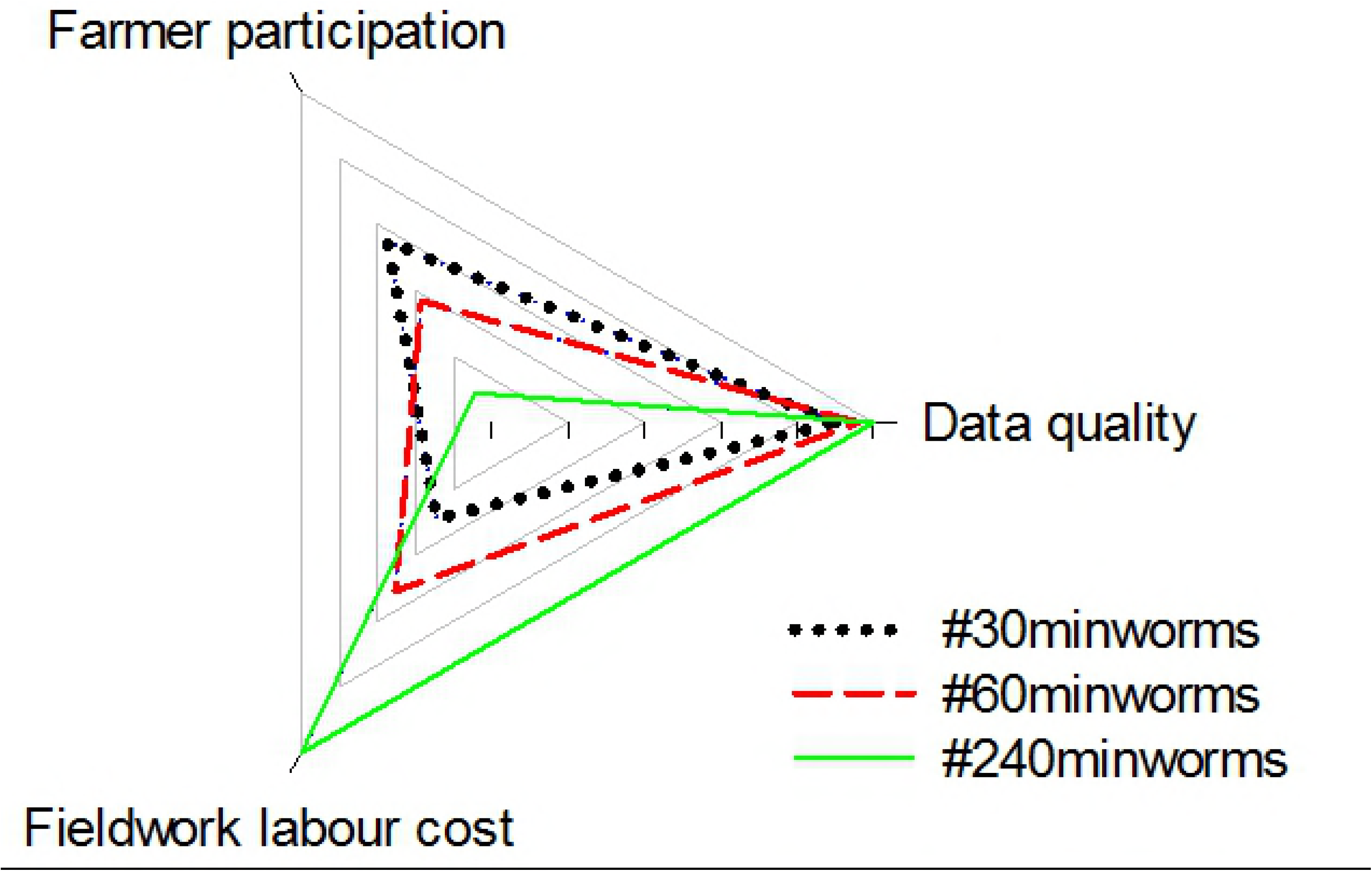
Trade-offs between earthworm fieldwork effort (30 – 240 mins) and data quality, farmer participation levels and labour costs.

## Discussion

The pilot #60minworms study effectively mobilised farmers to reach the target of 100 fields (Fig. 1). It was hypothesised that the workshops and newsletters would lead to the highest recruitment and participation rates due to a direct interaction and targeted approach (requiring a high time and cost), but posed a risk of location bias i.e. small geographic area monitoring. However, these channels had no impact on participation. Twitter, Farmers Weekly and The Farmers Forum were the most effective channels for recruitment. Twitter and Farmers Weekly recruits had exceptional participation and engagement rates, demonstrating the potential importance of these media channels to achieving soil security in agriculture. A high interest in community science was identified at the #60minworms workshop, with participants placing high value on others’ results, data collection abilities and motivations for sampling (Fig. 2), which would likely explain the impact of e.g. Twitter and Farmers Weekly over that of the isolated workshops and newsletters; with a further benefit of the wide geographic survey spread (Fig. 3). The community concept is further corroborated by the primary application of monitoring being to compare results within and between farms (64 %), and a high (87 %) interest in annual earthworm results from scientific national capability field trials e.g. Broadbalk indicating the potential to amplify both spatial and temporal soil health monitoring over and above what is achievable by these groups individually. Future developments that prioritize quick assessment protocols to enhance participation rates (farmers and number of fields), such as a #30minworms survey (Fig. 4) would likely be the most useful to farmers, as most participants (57 %) would change their soil management practices as a result earthworm monitoring results. This is in agreement with the ‘monitoring effect’, which is a confounding factor for gauging biodiversity[5], but is aligned with the DEFRA aspiration of sustainable soils by 2030. The absence of interest in regional data agrees with the primary interest in soil management (Fig. 2), and may explain the low participation rates by farmers in ecological earthworm surveys to date. At a national scale, £14 million pounds per #30minworms survey could be saved by mobilising farmers; demonstrating the potential high value of farmer input to achieving sustainable farmland soil policy.

The #60minworms method is a protocol validated for farmer applications, with feedback indicating high levels of trust, value and satisfaction by the participants (Fig.s 2, 3). There were no indications of significant sampling bias or problems in method compliance, however a key training need in earthworm identification skills was identified e.g. 46 % participants were not confident in their earthworm adult/juvenile separation and identification skills, but a significant interest in gaining this skill (Fig. 1). Farmer feedback led to modifications and improvement to the methodology and results presentation (SI booklets).

The findings from the #60minworm survey showed that earthworms are ubiquitous in UK farmland, with 100 % presence recorded on the majority (77 %) fields. The majority of these fields are managed under conventional agriculture (i.e. pesticides and inorganic fertilisers are used), and intensive cultivations have dominated crop establishment practices in England[15]. There was a significant (*p<* 0.05) impact of tillage on all parameters except endogeic earthworm presence (SI Fig. S1, Table S4). The survey revealed that there were no sightings of epigeic and anecic earthworm species, which are the two most sensitive ecological groups to tillage[13], on 21 % and 16 % fields respectively, and they were rare (≤10 % presence) on a further 8 % and 11 % fields (SI Table S5a-c). This is a cause for concern given the slow earthworm population recovery rates under changed management practices [33], and slow anecic earthworm reproduction rates, for example 8 cocoons per earthworm per year, with a 60 week development time [34]. No earthworm hotspots were detected in almost half (46 %) fields, where ≥ 16 worms per pit are linked to significant benefits in plant productivity (although this is highly dependent on a number of factors so does not have a strong interpretative value)[6]. At these measured on-farm population levels, these data indicate the majority of UK farmland soils have satisfactory earthworm biodiversity, but there is potential to increase the presence of these ecosystem engineers to better support both food security, but also wider earthworm-mediated ecosystem services such as native wildlife prey, soil aggregation and water infiltration; associated with soil security.

The aim of #60minworms was to indicate farmland soils at risk of over-cultivation through the absence/rarity of epigeic and anecic earthworms that have well known sensitivity to tillage. Here the ‘traffic light’ for results interpretation here was ranked as useful (36 %), but has an escalating error in categorizing earthworms at ≤10 sample pits, which could hinder participation whilst increase costs of monitoring (Fig. 4). Simplification is needed for a #30minworms survey, for example simply a ‘sub-optimal’ or ‘satisfactory’ score, the former indicated by < 20 % (b) epigeic, (c) endogeic and (d) anecic earthworm (or midden/vertical burrow) presence), would mitigate the problem of ‘false-negatives’ as both absent and rare (≤10 % presence) are within this ‘sub-optimal’ category (SI Table S5b,c). To aid the identification of exceptional earthworm populations for case-studies of soil management practices; Gold (100 %), Silver (≥80 %) and Bronze (≥60 %) ecological group presence could be used; of which 15 % of fields in this survey would have achieved a Gold or Silver rating. An alternative metric is the proposed soil health scorecard, using an identical size soil pit and hand-sorting, but at a sampling intensity of one pit per field and earthworm thresholds derived from Brazilian cropping systems [25]. In terms of quality control, there is a high labour cost (doubling of the hand-sorting assessment to 10-minutes for accuracy to improve the detection of juvenile worms), although a correction factor of 1.6 worms pit^-1^ could be used; the analysis may require five soil pits to provide a robust earthworm number estimate (SI Table S6) and this is a parameter with high annual variability (Table S1). The interpretation of ‘earthworm numbers’ is unclear, for example, earthworm numbers are linked to benefits in plant productivity, but this impact depends on soil texture, crop type and fertilisation regime [6], confounding the interpretative power of this parameter. A total of 20 % fields were identified as ‘depleted’ in earthworm numbers (Table 2), as this metric is primarily influenced by juvenile and endogeic earthworm abundance. In comparison, 42 % fields were depleted in adult ecological groups (principally epigeic and anecic earthworms with known vulnerability to tillage; good sources of food for native wildlife and roles in litter cycling and water drainage). This would explain why there is a limited concomitant relationship between the detection of ‘depleted’ fields using these interpretation schemes (Table 2). This could impact the ‘usefulness’ of earthworm data to farmers when interpretations of their fields significantly differ between scientists.

General strategies to increase the presence of earthworms would be to reduce tillage frequency and intensity (SI Fig. S1), however the impact of soil management activities is subject to local conditions (SI Table S3), and monitoring is an essential component to realising soil health in practice. One strategy that provides little benefit to earthworm populations is organic matter management (SI Fig. S2, Table S4). Three types of organic matter management were recorded, with straw retention or manuring having no significant (*p >* 0.05) impact on the presence of the ecological groups. However, cover cropping significantly (*p <* 0.05) increased the presence of anecic earthworms only (SI Fig. S2, Table S4). There was little evidence for organic matter management mitigating tillage impacts on earthworm populations. Identifying ‘at risk’ fields (up to 42 % fields in this survey), through the absence/rarity of epigeic and anecic earthworms, provides, for the first time, the opportunity for management intervention strategies to mitigate the effects of over-cultivation and support the DEFRA policy aspiration of sustainable soils by 2030.

## Acknowledgements

I’d like to thank the #60minworms participants for their invaluable inputs.

## Supporting information

**Supplementary Table S1.** Survey analysis using the hand-sorting data from multiple annual assessments on field trials managed under different organic matter rates and types. Despite large fluctuations in earthworm numbers, there was a consistent community structure.

**Supplementary Table S2.** Limited seasonal variation in earthworm community structures was detected on the AHDB Strategic Farm East in Autumn 2017 and Spring 2018 (n = 20 pits per field)

**Supplementary Table S3.** Field characteristics of the top and bottom 10 fields in the #60minworms survey.

**Supplementary Table S4.** P values from one-way ANOVA analyses of the #60minworms data set showing the significance of tillage on all parameters except endogeic presence. In comparison organic matter management practices of straw retention, cover cropping or manuring had little significant impact on earthworm parameters, with only cover cropping having a significant impact on anecic earthworm presence.

**Supplementary Tables S5a-c.** (a) The percentage of fields under earthworm ecological group presence categories, where no sightings are 0 % and may indicate a local extinction; and a likely presence is > 66 %, indicating there is good evidence for their presence based on 10 soil pits. (b) Fields with a sub-optimal ≤10 % presence (absent, rare) presence of earthworm ecological groups. (c) The percentage of fields under earthworm ecological group presence categories, where no sightings are 0 % and may indicate a local extinction; and a likely presence is > 66 %, indicating there is good evidence for their presence based on 5 soil pits.

**Supplementary Tables S6.** The field interpretation of earthworm counts at five pits compared to 10 pits is similar. However, there is high uncertainty at a low sampling intensity (one sample pit per field) as most fields (68 – 86 %) contain at least one pit (out of 10 pits) at each of the earthworm categories. This indicates that there is a considerable risk in over- estimating sub-optimal earthworm populations.

**Fig. S1.** The #60minworm survey results showed a negative impact (p < 0.05*) of tillage on earthworm presence (a, b, d, e) and numbers (f) (except endogeic presence).

**Fig S2.** The #60minworm survey found no significant (*p >* 0.05) impacts from straw retention or manuring management practices. Cover cropping had no significant (*p >* 0.05) impact on epigeic or endogeic earthworm presence, but a beneficial impact (*p <* 0.05*) on anecic earthworm presence.

**Supporting information S1 Booklet.** #60minworms Pilot study booklet, AHDB ‘How to count worm’ factsheets and new #30minworms booklet

